# Deletion of Autophagy gene *ATG1* and F-box motif encoding gene *YDR131C* together leads to Synthetic Growth Defects and Flocculation behaviour in *Saccharomyces cerevisiae*

**DOI:** 10.1101/2020.03.03.974519

**Authors:** Heena Shoket, Sadia Parvez, Meenu Sharma, Monika Pandita, Vishali Sharma, Prabhat Kumar, Narendra K Bairwa

## Abstract

F-box motif encoding *YDR131C* is functionally uncharacterized gene which forms the complex with the SCF-E3 ligase. The F-box motif containing proteins are involved in substrate recruitment for the ubiquitination and subsequent degradation through 26S proteasome. Autophagy gene, *ATG1 (ULK1*in human) is a well conserved serine-threonine kinase, required for vesicle formation and cytoplasm to vacuole targeting pathway. Atg1p forms the complex with Atg13p and Atg17p during autophagy. The understanding of crosstalk between ubiquitin and autophagy pathways is crucial for synthetic lethality screen and drug targeting. Here we have conducted the study for genetic interaction between uncharacterized *YDR131C* and *ATG*1 gene representing both specific and non-specific protein degradation pathways. The single and double gene knockout strains of *YDR131C*and *ATG1* genes were constructed in the BY4741 genetic background and analysed for growth fitness. The strains were also evaluated for cellular growth response in presence of hydroxyurea (HU), methyl methane sulfonate (MMS), and hydrogen peroxide (H_2_O_2_) stress causing agents by spot assay. The *ydr131cΔatg1Δ* showed the synthetic growth defect phenotype with floc formation in rich medium which showed floc disruption in presence of EDTA. The *ydr131cΔatg1Δ* cells showed the sensitivity to stress agents HU, MMS, and H_2_O_2_ when compared with *ydr131cΔ, atg1Δ*, and WT cells.. Based on the observations, we report that *YDR131C* and *ATG1* functions in parallel pathways for growth fitness and cellular growth response to stress agents. Interestingly this study also revealed the crosstalk between ubiquitination and autophagy pathways. The defects in both the pathways could lead to synthetic growth defects which may have implication for the precision medicine initiatives.

## Introduction

Ubiquitin proteasome pathway is very specific for degradation of smaller and specific poly-ubiquitinated proteins whereas non-specific and larger misfolded proteins requires autophagy pathway.Autophagy is highly conserved pathway for degradation of large protein complexes which include organelle material through lysosomal hydrolytic enzymes (*Budovskaya et al.* 2004). The autophagy pathway is well characterised which regulates various physiological processes and maintains the cellular homeostasis. The starvation and endoplasmic reticulum stress leads to induction of autophagy which involves formation of double membrane structure, known as auto-phagosome. Auto-phagosome encapsulates cytosolic components (organelles or proteins) to be fused with lysosome for degradation through hydrolases (Takeshige *et al.* 1992). In *Saccharomyces cerevisiae*, approximately 32 autophagy related genes (*ATG*) have been identified and one of these genes, *ATG1*, codes for cytosolic serine-threonine protein kinase, conserved from yeast to humans (Suzuki and Ohsumi 2007).The Atg1p is crucial for vesical formation and cytoplasm to vacuole targeting of the substrate (Matsuura *et al.* 1997; *Straub et al. 1997*). The Atg1p forms the complex with Atg13p and Atg17p for kinase activity during autophagy and is crucial for initiation of autophagy (Kabeya *et al.* 2005). It is also required for the localisation of the autophagy proteins such as Atg23p (Reggiori *et al.* 2004). Besides its role in autophagy as kinase factor, it is also function as structural protein which is involved in the PAS organization and assembly (Abeliovich *et al.* 2003). *S. cerevisiae* with null mutant of *ATG1* gene showed viability in nutrient rich medium (Giaever *et al.* 2002) and defects in autophagy (Matsuura *et al.* 1997), mitophagy (Zhang *et al.* 2007), pexophagy (*Hutchins et al. 1999*), cell death (Lisa-Santamaria *et al.* 2012), cytosol to vacuole transport (*Suzuki et al.* 2010), and sporulation. The *ATG1* is well conserved and its homologs have been reported in *C.elegans* (UNC-51) (Matsuura *et al.* 1997) and human (*ULK1*) (Kuroyanagi *et al.* 1998). The high throughput studies have reported many genes showing genetic interactions with *ATG1* (https://www.yeastgenome.org/locus/S000003148/interaction) involved in various biological processes. The genetic interaction of *ATG1* with mitochondrial protein encoding gene *AIM25* (Aguilar-Lopez *et al.* 2016) involved in maintaining the integrity of mitochondrial network and histone deacetylase, *HDA1* (Robert *et al.* 2011) and *RPD3* showed synthetic growth defects. The Rpd3 histone deacetylase regulates the transcriptional silencing and autophagy by regulating the chromatin remodelling activity (Robert *et al.* 2011).

F-box proteins are component of the SCF E3 ubiquitin ligase complex which recognizes the phosphorylated target proteins for ubiquitination and subsequent degradation by 26S proteasome. *YDR131C* of *S.cerevisiae* is non-essential gene, encodes for F-box motif protein (*Giaever et al.* 2002). The over expression of it leads to resistance against methyl mercury chloride in the W303 genetic background (Hwang *et al.* 2006). The null mutant of *ydr131cΔ* showed sensitivity to antimicrobial peptide, lacto-ferricin amino acids 1-11 (Lis *et al.* 2013) and methyl methane sulphonate (MMS) (Unpublished data from our lab). Genetic interactions between *YDR131C* with *ATG* genes have not been reported so far. Here, we report the genetic interaction between *YDR131C* and *ATG1* genes which regulates growth fitness, flocculation behaviour, and cellular response to genotoxic stress agents.

## Materials and Methods

### Yeast strain, plasmids, and growth conditions

*Saccharomyces cerevisiae* strain BY4741 (*MATa his3Δ1 leu2Δ0 met15Δ0 ura3Δ0*) was used for construction of single and double gene deletions (Table 1). Plasmids, pFA6a-KanMX6, and pFA6a-His3MX6 (Table 2) was used for amplification of the deletion cassettes for knockout of the *YDR131C* and *ATG1* ORFs. For ORF deletion PCR primers (Table 3) and method mentioned in (Longtine *et al.* 1998) was adopted.

**Table 1:** List of Yeast strains and their genotype used in this study

| S.no. | Strain | Genotype | Source |
| --- | --- | --- | --- |
| 1 | BY4741<br>(WT) | *MATa his3Δ1 leu2Δ0 met15Δ0 ura3Δ0* | Dr. Deepak Sharma, IMTECH, India |
| 2 | HS1 | *ydr131cΔ::KanMX* | This study |
| 3 | HNK1 | *atg1Δ::KanMX* | Dr. Ravi Manjithya, JNCASR, India |
| 4 | HNK2 | *ydr131cΔ::His3MX, atg1Δ::KanMX* | This study |

**Table 2:** List of plasmids used in this study

| S.no. | Plasmid Name | Deletion cassette | PCR product size | Selection Media |
| --- | --- | --- | --- | --- |
| 1 | pFA6a- KanMX6 | KanMX6 | 1559 bp | YPD + G418 |
| 2 | pFA6a- His3MX6 | His3M X6 | 1403 bp | SC His - |

**Table 3:**
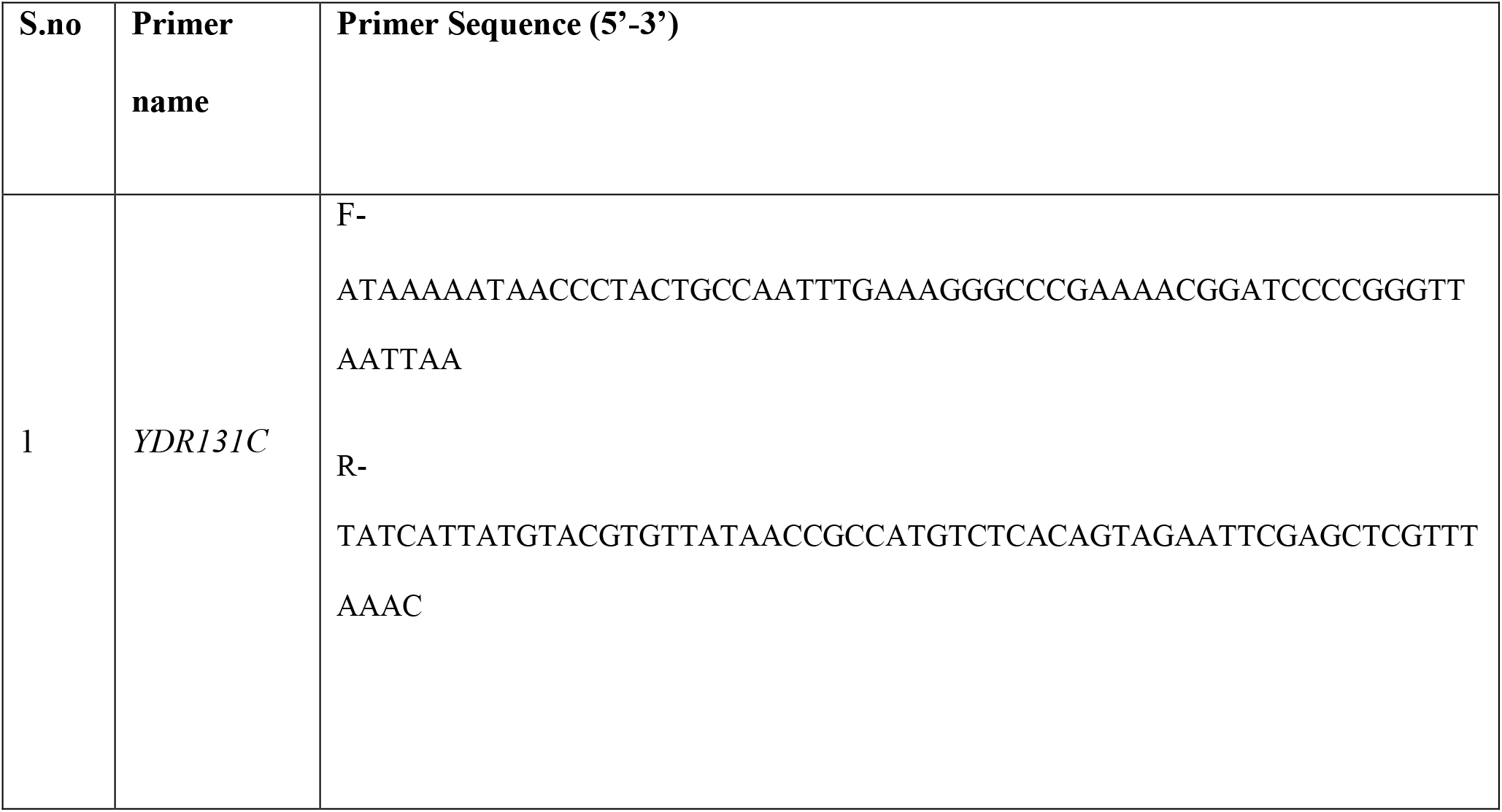
List of primers used in this study

### Growth Assay and Microscopy

Comparative growth assessment of WT, *ydr131cΔ, atg1Δ* and *ydr131cΔatg1Δ* was carried out both on solid and liquid media as mentioned in the (Sharma *et al.* 2019) and (Salari and Salari 2017). For comparative growth assessment on solid media each strains was streaked on YPD + agar plates and incubated for 36-48 hrs at 30° C. For comparative growth kinetics of strains in the liquid media, optical density measurement at 600nm for 14 hrs was carried out. For Comparative morphological assessment, an aliquot of each strain from liquid media was observed under Leica DM3000 microscope at 100X magnification.

### Growth in YPD medium and Flocculation behaviour

The WT, *ydr131cΔ, atg1Δ* and *ydr131cΔatg1Δ* cells grown in YPD medium overnight and observed for floc formation visually and using microscope. The dependence of the floc formation on EDTA was tested by adding the 30mM of EDTA (chelating agent) to growth medium (Stewart 2018).

### Spot Assay

To compare the cellular growth response of WT, *ydr131cΔ, atg1Δ*,and *ydr131cΔatg1Δ* cells in presence of stress causing agents spot assay was performed as mentioned in (*Sharma et al.* 2019). Briefly Wild type (BY4741) and deletion derivatives (WT, *ydr131cΔ, atg1Δ* and *ydr131cΔatg1Δ*) were grown to log phase (OD^600^ 0.8-1.0) and equal number of cell were serially diluted. From each dilution a 3µl aliquot was spotted onto agar plates containing YPD, YPD + stress causing agents such as a hydroxyurea (200mM), MMS (0.035%) and hydrogen peroxide (4mM). The plates were incubated at 30°C for 2-3 days and imaged.

### Fluorescence Microscopy for Cell Wall and Nuclear status

To compare the cell wall status of WT, *ydr131cΔ, atg1Δ*, and *ydr131cΔatg1Δ* cells, Calcofluor white staining assay described in (Pringle 1991; Preechasuth *et al.* 2015; *Sharma et al.* 2019) was adopted. Briefly, WT and mutant strains were grown overnight at 30ºC and next day transferred to fresh culture in 1:10 ratio. Cells were grown to log phase and collected by centrifugation. Further, cells were suspended in 100µl of solution containing Calcofluor white (50 μg/ml solution) fluorescent dye. For nuclear status by DAPI staining log phase grown cells were harvested by centrifugation, washed with distilled water, and suspended in 1X Phosphate Buffer Saline (PBS). Further, fixation was done by adding 70% ethanol for 10 minutes and centrifuged for 2.5 minutes at 2500 rpm. DAPI stain (1mg/ml stock) to final concentration 2.5µg/ml was added and incubated for 5 minutes at room temperature and visualized under UV light of fluorescent microscope with 100X magnification using Leica DM3000 fluorescence microscope.

## Results

### Loss of *YDR131C* and *ATG1* together leads to synthetic growth defect and Flocculation behaviour

The null mutant of both the gene *YDR131C* and *ATG1* have been reported to be viable. The comparative growth assessment of both *ydr131c*Δ and *atg1Δ* strains was similar to the growth of WT cells on solid and liquid media (Figure 1A, B). However *ydr131cΔ atg1Δ* cells showed slow growth phenotype when compared with WT*, ydr131cΔ*, and *atg1Δ* mutants indicating synthetic growth defect phenotype. The microscopy analysis of the WT, *ydr131cΔ, atg1Δ* and *ydr131cΔ atg1Δ* cells when grown in rich media (YPD), the *ydr131cΔatg1Δ* cells showed the floc formation during growth kinetics (Figure 1C, 2A,B). Flocculation is cell surface characteristics and a process in which the cells have the ability to form clumps. The flocculation is regulated by genetic, physiological, and environmental factors. Further when cells *ydr131cΔatg1Δ* were grown in YPD media supplemented with 30mM EDTA, the floc formation behaviour was reduced (Figure 2C). This indicated, the floc formation of the *ydr131cΔatg1Δ* was EDTA dependent.

**Figure 1.**
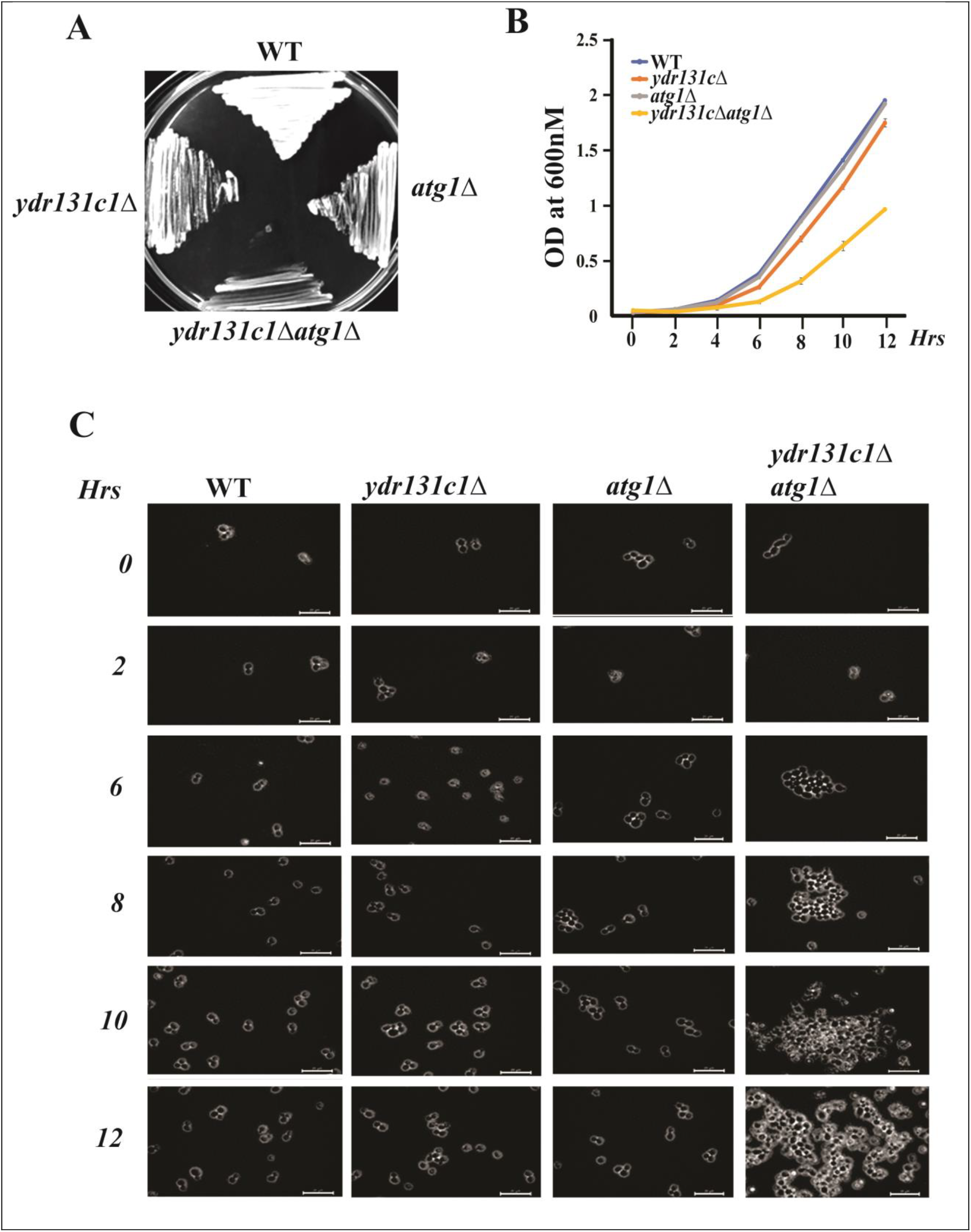
Comparative assessment of growth and morphological characteristics of WT, *ydr131c∆, atg1∆ and ydr131c∆atg1∆* cells. **A.** Comparative growth on YPD solid medium **B.** Growth kinetics of WT, *ydr131c∆, atg1∆, and ydr131c∆atg1∆* strains *YPD* liquid medium. The error bars represent the standard deviation of three independent experiments **C.** Phase contrast images of WT, *ydr131c∆, atg1∆, and ydr131c∆atg1∆* strains taken during analysis of growth kinetics at indicated time point. Images were acquired at 100X magnification using Leica DM3000 microscope. The *ydr131c∆atg1∆* cells showed floc formation behaviour.

**Figure 2.**
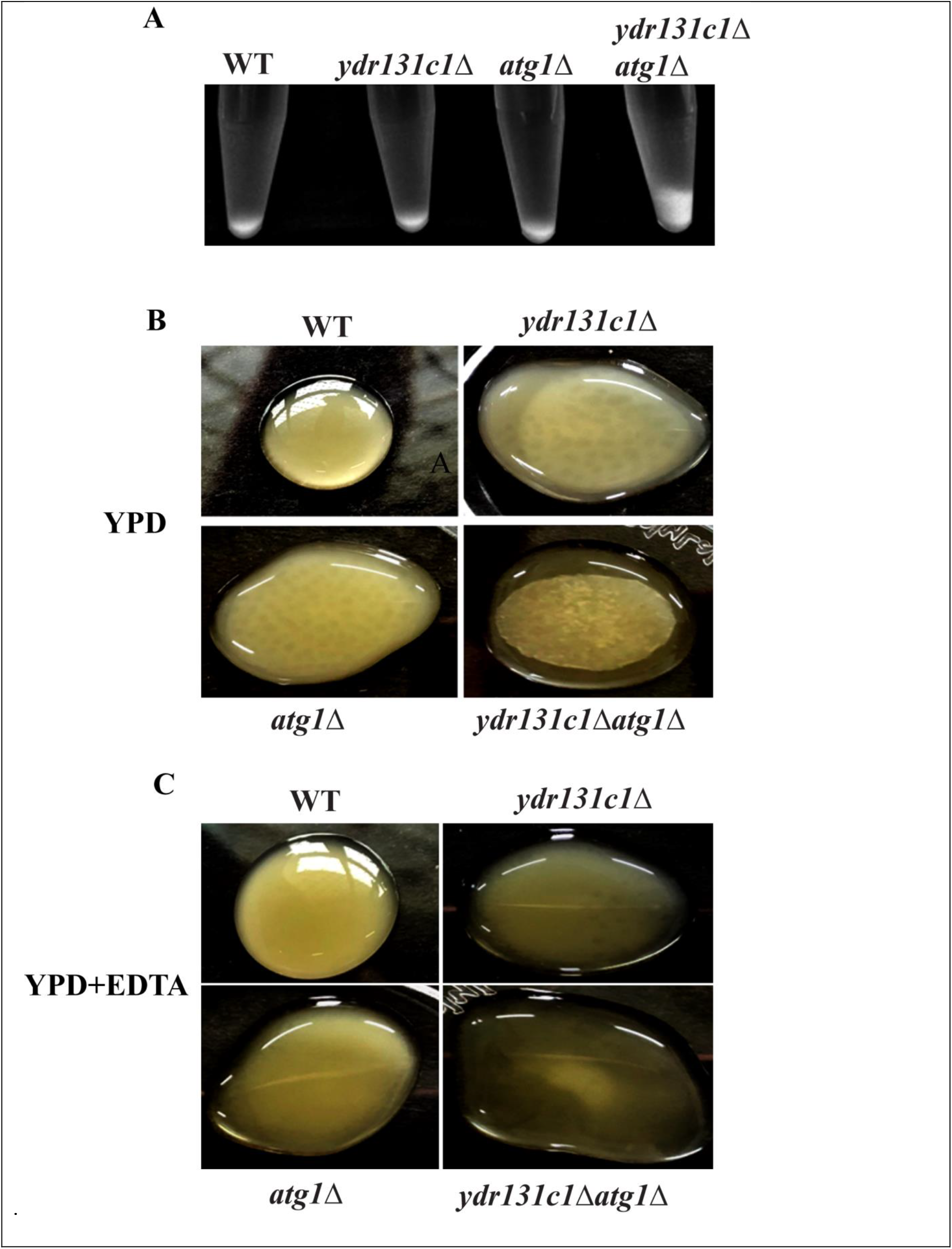
Comparative assessment of the floc formation in WT, *ydr131c∆, atg1∆, ydr131c∆atg1∆* strains. ***A.*** Overnight grown WT, *ydr131c∆, atg1∆, ydr131c∆atg1∆* strains were poured in individual tubes and allowed to settle for 30 mins and then imaged. The *ydr131c∆atg1∆*cells showed the diffused settlement B. Overnight grown cultures was gently poured in the centre of petri dish and allowed to settle, ydr131c*∆atg1∆* cells formed th**e** floc. **C.** WT, *ydr131c∆, atg1∆*, and *ydr131c∆atg1∆* cells grown in presence of 30mM EDTA overnight showed disappearance of floc formation.

### Loss of *YDR131C* and *ATG1* together does not affect chitin distribution

Calcofluor white is a flourochrome stain, which is specific for the cell wall component chitin and has been used for detection of cell wall defect (DE GROOT et al. 2001). The Calcofluor white staining of WT, *ydr131cΔ, atg1Δ*, and *ydr131cΔatg1Δ* showed clear septa formation however *ydr131cΔatg1Δ* showed aggregated cells and altered bud morphology (Figure 3).

**Figure 3.**
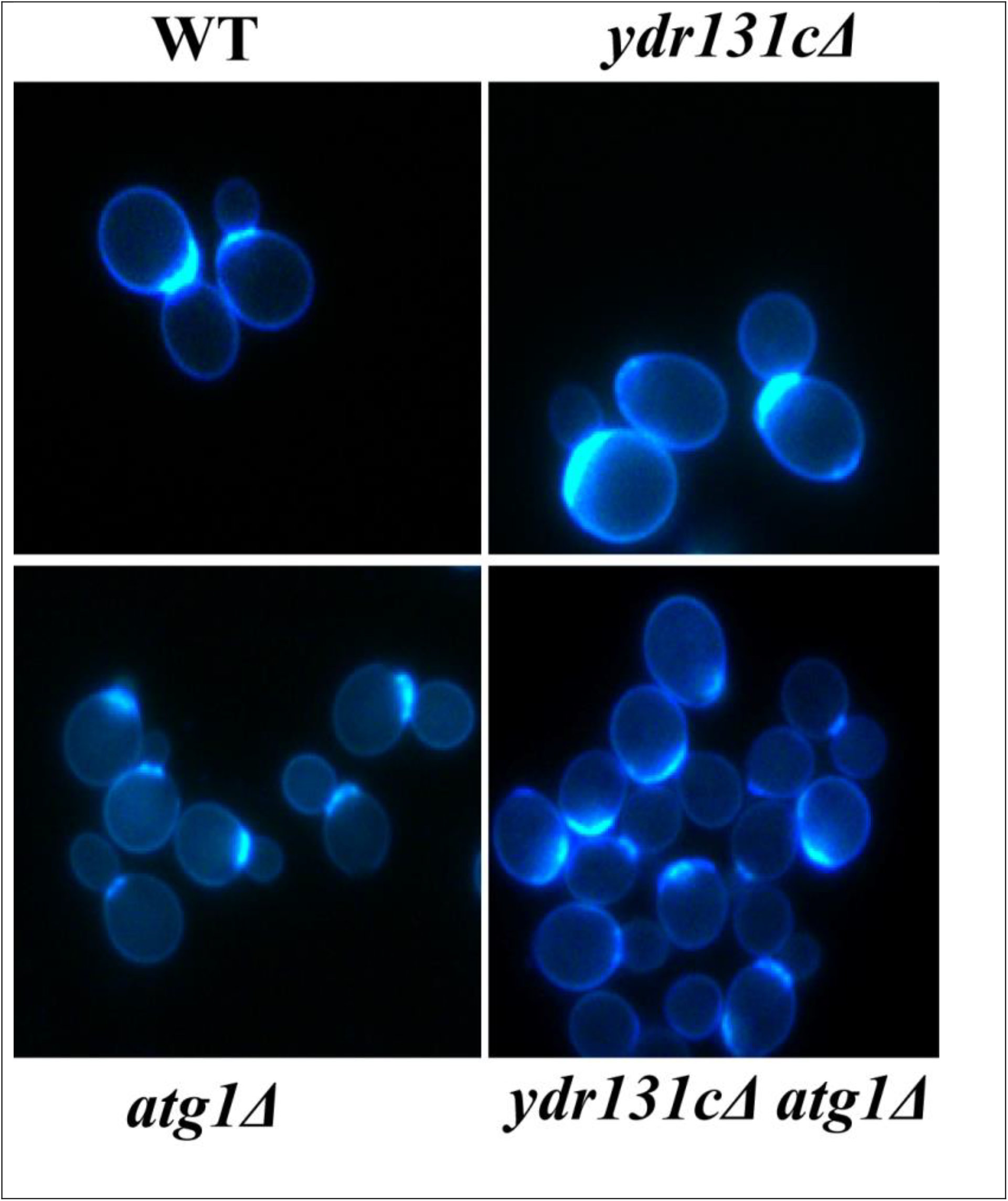
Comparative assessment of chitin distribution in WT, *ydr131c∆, atg1∆*, and *ydr131c∆atg1*∆ by Calcofluor white staining. *The ydr131c∆atg1*∆ showed no altered chitin distribution in comparison to WT, *ydr131c∆*, and *atg1∆* cells.

### Loss of *YDR131C* and *ATG1* together leads to sensitivity to genotoxic and oxidative stress

Genotoxic stress agents, hydroxyurea (HU) and methyl methane sulphonate (MMS) act as DNA damaging agent whereas hydrogen peroxide leads to oxidative stress, respectively. The cellular growth response of WT, *ydr131cΔ, atg1Δ* and *ydr131cΔatg1Δ* by spot assay in presence of 200mMHU, 0.035%MMS and 4mM H_2_O_2_ indicated, extremely slow growth of *ydr131cΔatg1Δ* cells when compared with the WT and *ydr131cΔ*, a*tg1Δ* (Figure 4 A,B,C).

**Figure 4.**
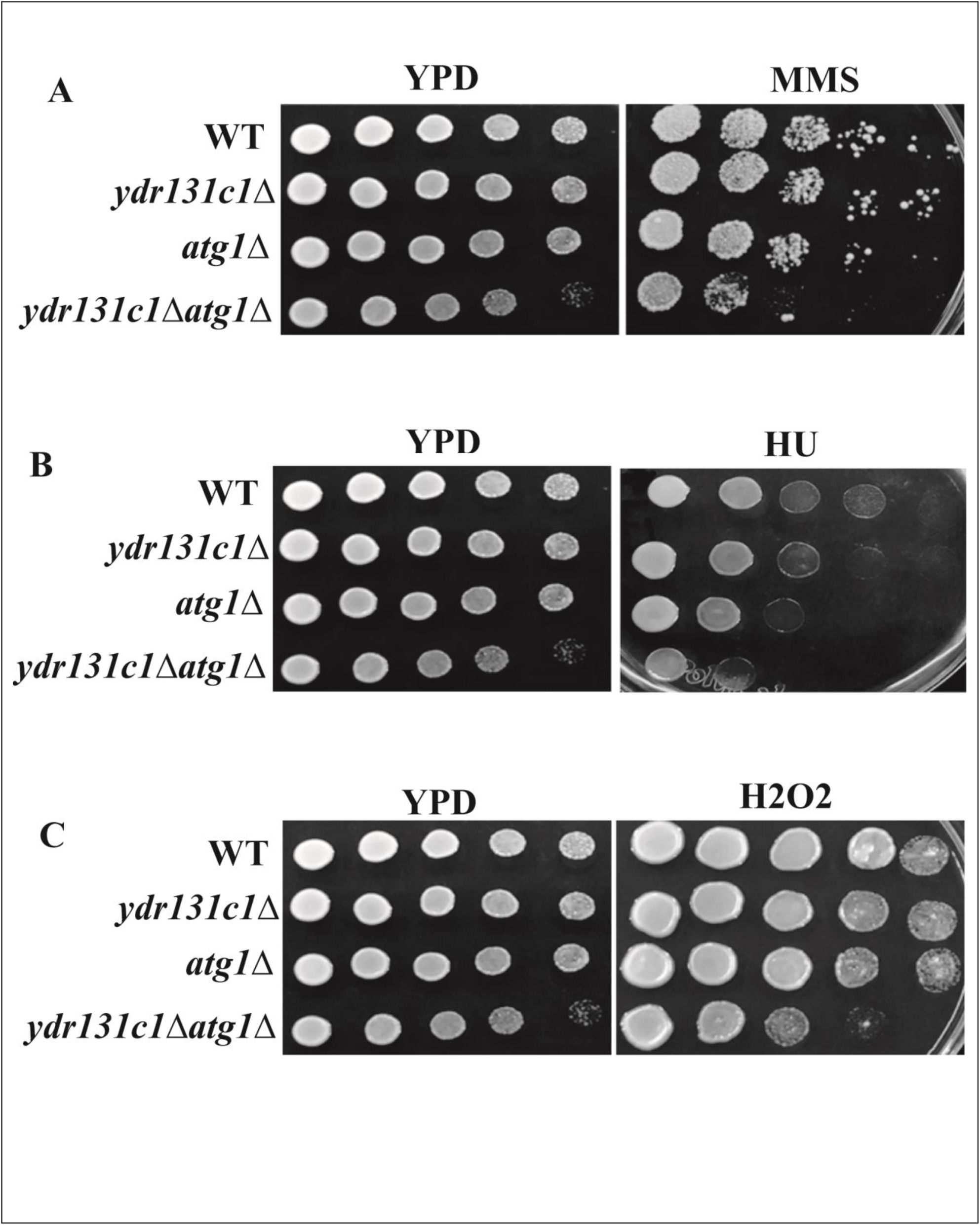
Comparative assessment of cellular growth response of WT, *ydr131c∆, atg1∆ and ydr131c∆atg1∆* cells to genotoxic and oxidative stress agents. Equal number of Log phase grown cells of each strain serially diluted and spotted on YPD agar plate and plates containing 0.035% methyl methane sulfonate, 200mM hydroxyurea and 4mM hydrogen peroxide. The *ydr131c∆atg1∆* cells showed sensitivity to the genotoxic and oxidative stress agents in comparison to WT, *ydr131c∆*, and *atg1∆* cells.

### Loss of *YDR131C* and *ATG1* together leads to altered nuclear morphology

DAPI stain is specific to the nuclear DNA which allows the visualization of the DNA in the form of single or defective nuclei depending upon the cells. The comparative analysis of nuclei in WT, *ydr131*cΔ*, atg1Δ*, and *ydr131cΔatg1Δ* of the log phase culture stained with DAPI showed compact nuclei in WT, *ydr131cΔ, atg1Δ* whereas *ydr131cΔatg1Δ* cells showed alerted nuclear morphology (Figure 5)

**Figure 5.**
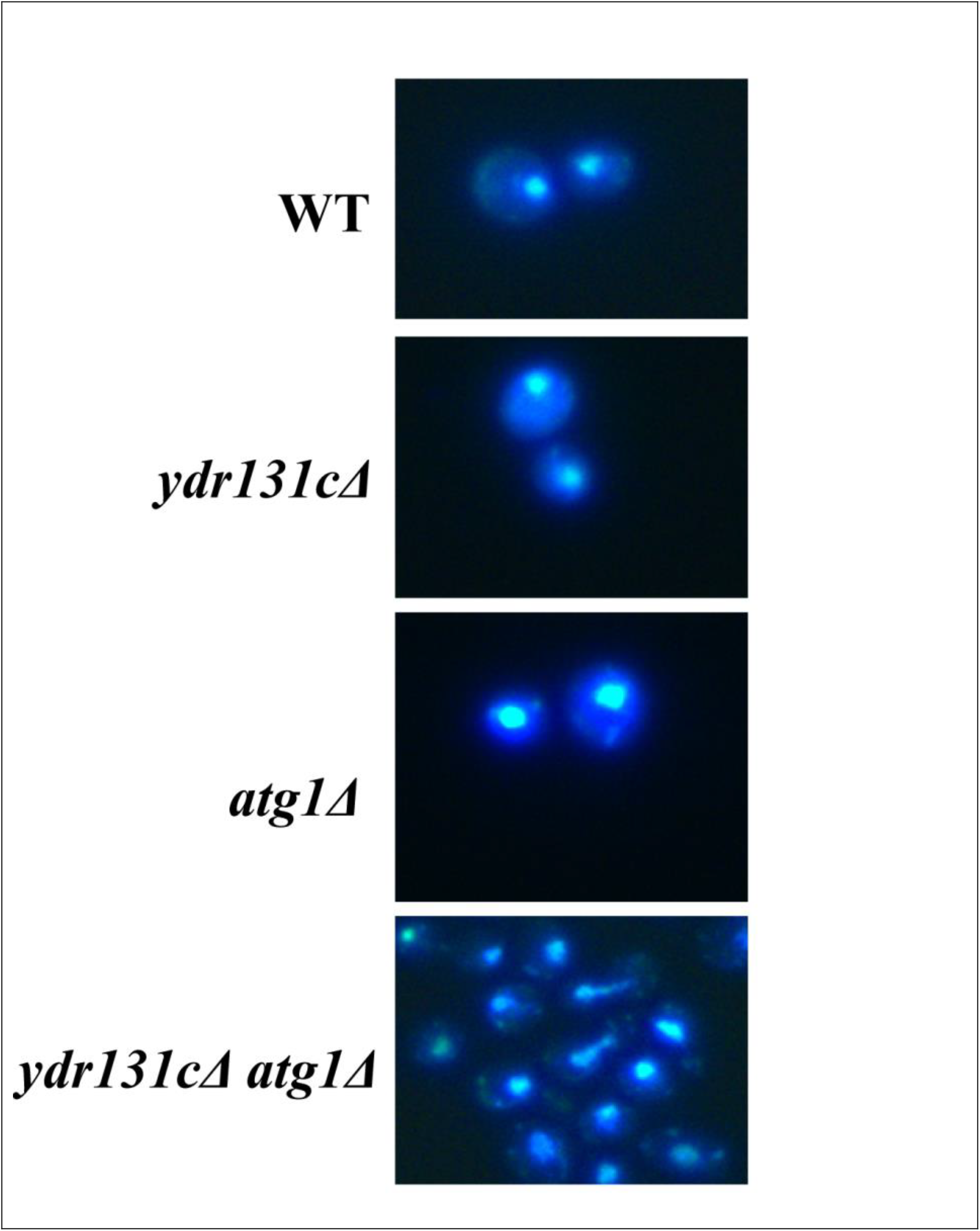
Comparative assessment of nuclear status in WT, *ydr131c∆, atg1∆, and ydr131c∆atg1∆* using DAPI (4′, 6-diamidino-2-phenylindole) staining and fluorescence microscopy. The *ydr131c∆atg1∆*cells exhibited diffused nucleus as compared to WT, *ydr131c∆* and *atg1∆*.

## Discussion

The novel binary interactions between gene pairs and their functional relevance are crucial for better understanding of wiring of the pathways in the cell system. The Skp1-Cul1-F-box (SCF) E3 ligase complex recruits the target proteins for ubiquitination by its one of subunit protein which contains F-box motif, for degradation by the 26S proteasome (Bai *et al.* 1996; Hershko and Ciechanover 1998). The Ydr131c is F-box protein whose target has not been reported so far (Kipreos and Pagano 2000) however overexpression of it lead to resistance to methylmercury (Hwang *et al.* 2006). Here we observed the null mutants of *YDR131C* showed sensitivity to genotoxic agents such as hydroxyurea and MMS. Autophagy gene *ATG1* is crucial for autophagy response, the lack of it leads to absence of apoptosis, autophagy, and pexophagy in the cells. The combined mutation in autophagy gene *ATG*1 and *YDR131*C showed synthetic growth defects and altered nuclear morphology suggesting that both the genes are required for growth fitness and cellular response to the hydroxyurea, MMS, and oxidative stress. In addition the flocculation behaviour observed with *ydr131cΔatg1Δ* cells is an interesting finding which needs to be investigated in the future. The investigated genetic interaction may have implication for synthetic lethal pair discovery for cancer, as there is high degree of functional homology of *ATG1* with mammalian *ULK1*. The mechanism of crosstalk between the specific protein degradation pathway mediated by *YDR131C* and non-specific proteins degradation pathway mediated by *ATG1* reported here would be investigated in future.

## Acknowledgment

The authors would like to thank Dr. Deepak Sharma, IMTECH, Dr. Ravi Manjithya, JNCASR, Dr. Jitendra Thakur, NIPGR, New Delhi, India for strains and plasmids. We also thank Dr. Preeti Sharma and Mr. Parvez S. Slathia at SMVDU for allowing the use of lab equipment facility.

## Compliance with Ethical Standards

Authors declares no conflict of interest.

## Ethical Approval

This article does not contain any studies with human participants performed by any of the authors.

## Funding information

The research work in the laboratory of N.K.B is supported by Ramalingaswami fellowship grant (BT/RLF/Re-entry/40/2012) from the Department of Biotechnology and SERB-DST, GOI grant number (EEQ/2017/0000087) and support from SMVDU, Jammu & Kashmir, India.

## Author’s contributions

NKB conceived and directed the study and wrote the paper with all the authors. HS, SP, MS, VS, MP, PK conducted the experiments. All the authors analysed the data, reviewed the results, and approved the final version of manuscript.

